# Peripheral Myelin Protein 22 Preferentially Partitions into Ordered Phase Membrane Domains

**DOI:** 10.1101/2020.01.28.923771

**Authors:** Justin T. Marinko, Geoffrey C. Li, Anne K. Kenworthy, Charles R. Sanders

## Abstract

The ordered environment of membrane rafts is thought to exclude many transmembrane proteins. Nevertheless, some multi-pass transmembrane proteins have been proposed to partition into ordered domains. Here, giant plasma membrane vesicles (GPMVs) were employed to quantitatively show that the tetraspan peripheral myelin protein 22 (PMP22) exhibits a pronounced preference for, promotes the formation of, and stabilizes ordered membrane domains. Neither S-palmitoylation of PMP22 nor its putative cholesterol binding motifs are required for partitioning to ordered domains. In contrast, disruption of its unusual first transmembrane helix (TM1) reduced ordered phase preference. Charcot-Marie-Tooth disease-causing mutations that destabilize PMP22 also reduced or eliminated this preference in favor of the disordered phase. These studies demonstrate PMP22’s ordered phase preference derives both from the distinctive properties of TM1 and global structural features associated with its transmembrane domain, providing a first glimpse at the structural factors that promote raft partitioning for multi-pass proteins.

**Significance Statement:** The preferential partitioning of single span membrane proteins for the ordered phase of ordered/disordered phase-separated membranes is now reasonably well understood, but little is known about this phase preferences of multi-pass membrane proteins. Here, it is shown that the disease-linked tetraspan integral membrane protein, PMP22, displays a pronounced preference to partition into the ordered phase, a preference that is reversed by disease mutations. This phase preference may be related to the role of PMP22 in cholesterol homeostasis in myelinating Schwann cells, a role that is also known to be disrupted under conditions of CMTD peripheral neuropathy caused by *pmp22* mutations.

## Introduction

Our current understanding of biological membranes has been shaped over the past 30 years by studies of membrane phase separation into ordered and disordered domains. Early on, these studies yielded the lipid raft hypothesis (*1, 2*) which was hotly debated in subsequent years (*3–8*). Based on a wealth of data it is now generally believed that phase separation does sometimes occur in sphingomyelin and cholesterol-rich membranes, such as in eukaryotic plasma membranes (PMs). While it is thought that phase-separated ordered are often small in size, transient, and similar to the adjacent disordered phase in lipid composition (*8–10*), there also appear to be certain native membranes that are so cholesterol and sphingolipid-rich that their physical properties are, to a significant degree, akin to those of ideal liquid-ordered (Lo) phase model membranes (*9, 11–13*)

Early studies of membrane protein association with raft-like ordered membrane domains was based largely on results involving the isolation of “detergent-resistant membranes” (*14–17*). More recent biophysical studies conducted in intact phase-separated membranes have confirmed that a number of single-pass transmembrane proteins do indeed have a preference to partition into ordered phase domains relative to conjoined disordered phase domains. Particularly important in this regard are studies that have employed “giant plasma membrane vesicles” (GPMVs), which can be formed from a variety of mammalian cell types (*12*). When GPMVs are isolated and then cooled, separation of microscopically-observable ordered and disordered membrane phases can occur, allowing for quantitative studies of protein partitioning between the two phases (*3, 12, 18-21*). GPMVs therefore provide facile experimental access to conditions in which large and stable ordered phase domains co-exist with disordered membranes (*18, 21, 22*). Groundbreaking studies of single span membrane proteins in this medium led to development of a convincing quantitative model describing the structural basis for why some proteins of this class preferentially partition into the ordered phase (*12, 20, 23-26*). Studies of the phase preferences of multi-pass membrane proteins remain at a much earlier stage of development. Here, we present the first example of a multi-pass membrane protein that exhibits a pronounced preference for the ordered phase in GPMVs—the tetraspan peripheral myelin protein 22 (PMP22).

The human PMP22 is a 160-residue protein containing four transmembrane helices and intracellular N- and C-termini (**Fig. 1A**). PMP22 appears to play multiple roles in myelinating Schwann cells and peripheral myelin (*27–31*) including a critical role in cholesterol homeostasis, particularly in trafficking of cholesterol through the secretory pathway to the membrane surface (*32*). This is especially important in Schwann cells given their specialized function as the factory for myelin production in the peripheral nervous system (PNS). Myelin membranes are unusually rich in both cholesterol and sphingolipids (*33, 34*) and are therefore highly ordered, as well suits their roles in providing electrical insulation and mechanical support to PNS axons. Mutations in the *pmp22* gene result in >70% of all cases of Charcot-Marie-Tooth disease (CMTD, prevalence: 1:2500) and related peripheral neuropathies (*28, 29, 35*). These closely related disorders are characterized by defective myelin membranes that contain altered cholesterol levels relative to healthy myelin (*36–38*).

**Figure 1.**
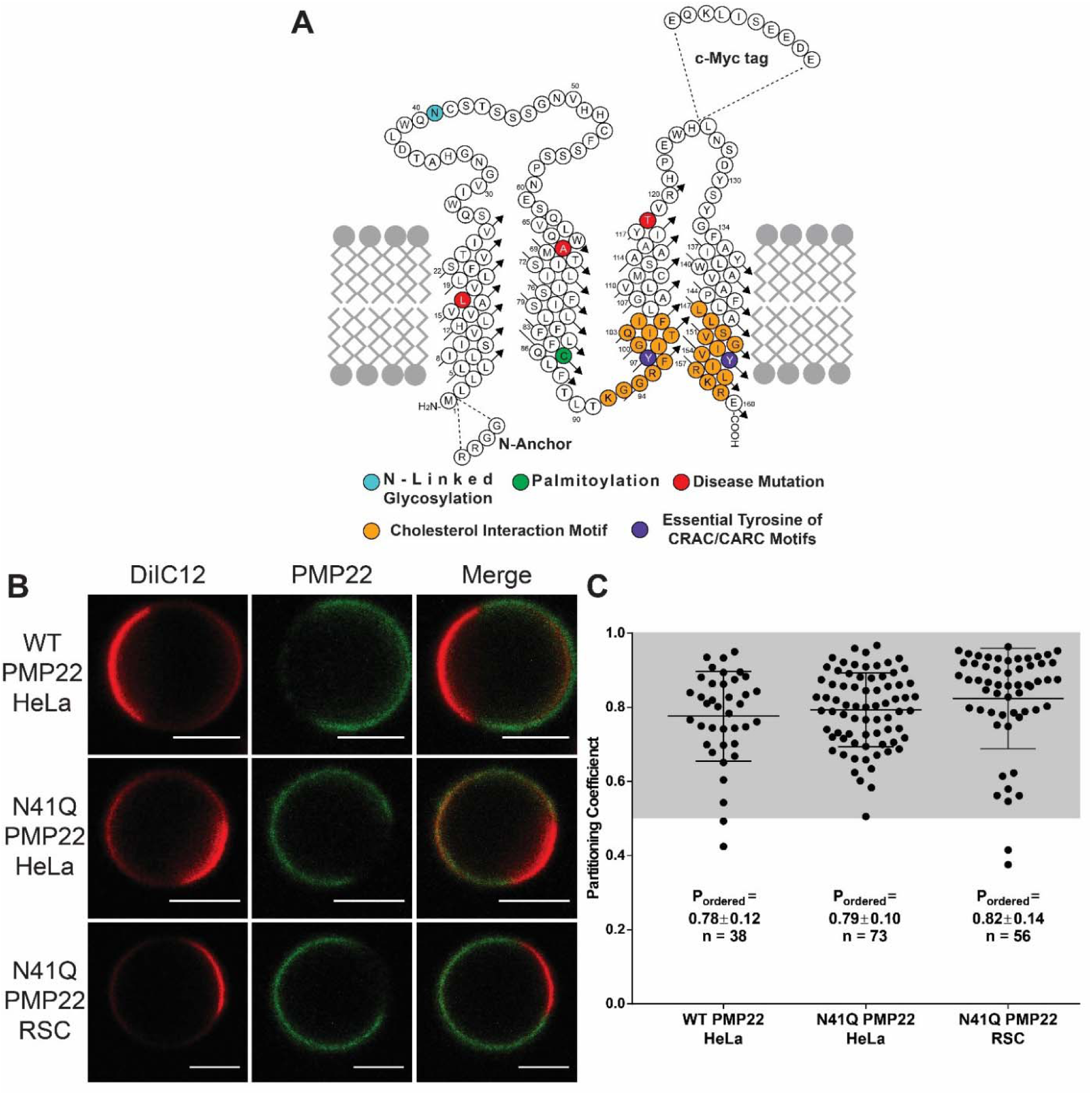
PMP22 partitions into ordered phase domains of GPMVs. **(A)** Cartoon topology map of PMP22 showing the locations of the myc-epitope tag, sites of post-translational modifications (cyan: glycosylation green: palmitoylation), cholesterol interaction motifs (orange) and their essential Tyr residues (purple), N-anchor insertion site, and the sites of the CMTD disease mutations examined in this study (L16P, A67T, T118M red). **(B)** Representative GPMVs containing WT or N41Q PMP22 derived either from HeLa cells or RSCs. Scale bar, 5 µm. The disordered phase marker is shown in red and PMP22 is shown in green. **(C)** Quantification of PMP22 partitioning coefficients from three independent biological experiments for each condition. Each point represents an individual GPMV. Mean ± standard deviation (SD) is reported and plotted on graph.

The involvement of PMP22 in cholesterol trafficking as part of the process of myelin membrane formation suggests that this protein might have an intrinsic affinity for ordered membrane domains. Indeed, it has previously been reported that PMP22 is found in “lipid raft” membranes isolated from neurons following application of classical detergent-extraction methods (*34, 39*). However, some of the methods used for preparing lipid rafts, including that used to identify PMP22 as a raft-associated protein, are thought to be artifact-prone (*6, 7, 14-16*). Nevertheless, the likely role of PMP22 in Schwann cell cholesterol homeostasis combined with its residence in cholesterol and sphingolipid-rich myelin membranes dictate that the hypothesis that PMP22 may preferentially partition into ordered phase membranes is compelling. This hypothesis is tested in this work.

## Results

### PMP22 preferentially partitions into ordered phase membrane domains of GPMVs

To examine the preference of PMP22 for the ordered versus the disordered phase in plasma membranes (PM), we expressed human PMP22 (**Fig. 1A**) in HeLa cells and then prepared GPMVs using established protocols (*21*). To render PMP22 easily immunodetectable, the c-myc tag was inserted into the second extracellular loop, a modification that has no effect on PMP22 trafficking or function (*40, 41*). PMP22 in GPMVs was visualized using confocal fluorescence microscopy using an anti-myc antibody conjugated to AlexaFluor 647 (AF647, green). Disordered phase domains within GPMVs were identified using the fluorescent carbocyanine lipid DiIC12 (red), which partitions preferentially into disordered membrane domains (*22*). **Fig. 1B** shows representative PMP22-containing GPMVs. Within each phase PMP22 was uniformly distributed and showed no tendency to concentrate at the boundary between the ordered and disordered domains (**Fig. 1B**).

In our experiments we noticed an increase in the amount of PMP22-containing GPMVs when preparations were performed using cells overexpressing the N41Q N-glycosylation deficient variant of PMP22 (**Fig. 1A**, cyan) versus the wild type (WT) protein. Eliminating glycosylation of PMP22 does not affect its function or turnover (*42, 43*). This observation is most likely due to an increased concentration of N41Q PMP22 at the PM compared to WT PMP22 (**Fig. S2**). Representative N41Q PMP22-containing GPMVs derived from both HeLa cells and primary rat Schwann cells (RSCs) are also shown in **Fig. 1B**. Both glycosylated and non-glycosylated PMP22 show a clear preference for ordered phase membrane domains, as evidenced by the lack of co-localization of the red DiIC12 and green AF647 channels. PMP22 shows a distinct preference for ordered membrane domains in GPMVs prepared from both HeLa cells and RSCs.

AF647 fluorescence intensity was quantified in ordered and disordered GPMV membrane domains to determine the relative concentration of PMP22 in each membrane phase. Following image quantitation, the partitioning coefficient (P_ordered_) of PMP22 was calculated, where P_ordered_ is: [PMP22]_ordered_/([PMP22]_ordered_ + [PMP22]_disordered_), and ranges from 0 to 1 with a value of 0.5 meaning the protein has equal affinity for both phases, while P_ordered_ > 0.5 means that the protein prefers the ordered phase, and P_ordered_ < 0.5 means that the protein prefers the disordered phase. For an example of GPMV analysis see **Fig. S1** (*25*).

GPMVs from HeLa cells showed clear phase separation at 20°C while those from RSCs exhibited phase separation at 15°C. Quantification of P_ordered_ for WT and N41Q PMP22 in GPMVs derived from HeLa cells as well as for N41Q PMP22 in GPMVs derived from RSCs is shown in **Fig. 1C**. Data was acquired from three independent biological experiments for each condition. In HeLa GPMVs, WT PMP22 showed a P_ordered_ of 0.78±0.12 (mean ± SD) and N41Q PMP22 showed a P_ordered_ of 0.79±0.10. N41Q PMP22-containing RSC GPMVs displayed a P_ordered_ of 0.82±0.14. As a control, we measured the phase partitioning in both HeLa cells and RSC GPMVs of the well-studied mEGFP-labeled form of the single pass membrane protein, linker for activated T Cells (tgLAT) (**Fig. S3**). The measured P_ordered_ values of 0.54±0.11 and 0.55±0.07 for tgLAT in GPMVs derived from Hela and RSCs respectively are in line with those reported in the literature using the same GPMV preparation in rat basal leukemia cells (*20, 24*). Additionally, we measured P_ordered_ for N41Q PMP22 in HeLa GPMVs using a different membrane phase marker, NBD-DSPE (**Fig. S4**), which identifies ordered membrane phase domains (*22*). Using this marker, we calculated a P_ordered_ of 0.82±0.06. These results quantitatively demonstrate that PMP22 has a significant preference to partition into ordered membrane domains in GPMVs from both model mammalian cell lines and primary Schwann cells. Additionally, this phase preference is not affected by N-linked glycosylation. There were no significant differences in WT versus N41Q PMP22 phase partitioning. In light of this results and because the level of surface expression for N41Q PMP22 made it easier to image, all subsequent experiments reported in this work utilized a N41Q PMP22 variant (hereto referred to as “PMP22” for the sake of simplicity). Additionally, because HeLa-derived GPMVs showed phase separation at a temperature closer to physiological levels and were easier to transfect compared to RSCs, all subsequent experiments utilized HeLa derived GPMVs.

### S-Palmitoylation of PMP22 is not a significant driver of its ordered phase preference

It was recently shown that PMP22 is palmitoylated at Cys85 (**Fig. 1A,** green) (*44*). In that study this post-translational addition of a saturated fatty acid group on the cytosolic side of the protein was shown not to affect PMP22 processing/trafficking but is important for modulating epithelial cell shape and motility. For single pass transmembrane proteins, it has been repeatedly shown that palmitoylation plays a significant role in mediating membrane phase partitioning; one study estimated that this modification contributes ∼0.5 kcal mol^-1^ free energy per chain in favor or ordered phase partitioning (*20, 24*). The removal of palmitoylation from tgLAT disrupts the ordered phase preference of that protein and causes it to equally prefer both membrane phases (*24*). We therefore tested to see if palmitoylation affects PMP22 partitioning into ordered membrane phases.

To eliminate the palmitoylation of PMP22 we mutated Cys85 to an Ala residue. This mutation did not affect the localization of PMP22 in HeLa cells (**Fig. S2**). We then measured the P_ordered_ for C85A PMP22 in GPMVs. As seen in **Fig. 2A-B** palmitoylation did not drastically affect ordered phase preference of PMP22. We determined a P_ordered_ for C85A PMP22 of 0.77±0.10. This value is almost identical to that reported for WT and N41Q PMP22 in **Fig. 1**. Because our method of GPMV preparation requires the use of the reducing agent dithiothreitol (DTT; 2 mM) we also set tested if PMP22 was still palmitoylated under these conditions. PMP22 transfected cells were incubated overnight with 100 mM of 17-octadecynoic acid (17-ODYA), a palmitic acid analogue containing an alkyne on the terminal carbon, which is known to be incorporated by thioesterases into palmitoylated proteins. Cells were then treated with or without 2 mM DTT for 90 min at 37°C, lysed, and PMP22 was immunoprecipitated. We then added a biotin handle to palmitoylated proteins using biotin azide and classical ‘click’ chemistry (*44*). Palmitoylated PMP22 was identified via western blotting using an anti-biotin antibody (**Fig. 2C).** Palmitoylation was quantified as the intensity of the anti-biotin band over the anti c-myc band and normalized to the amount of palmitoylation found in N41Q PMP22 samples without DTT treatment (**Fig. 2D**). As seen in **Fig. 2C-D**, treatment of HeLa cells for 90 minutes with 2 mM DTT did not affect PMP22 palmitoylation. As expected, C85A PMP22 was not palmitoylated in these experiments. These results show that S-palmitoylation of PMP22 is not a significant driver of ordered phase preference.

**Figure 2.**
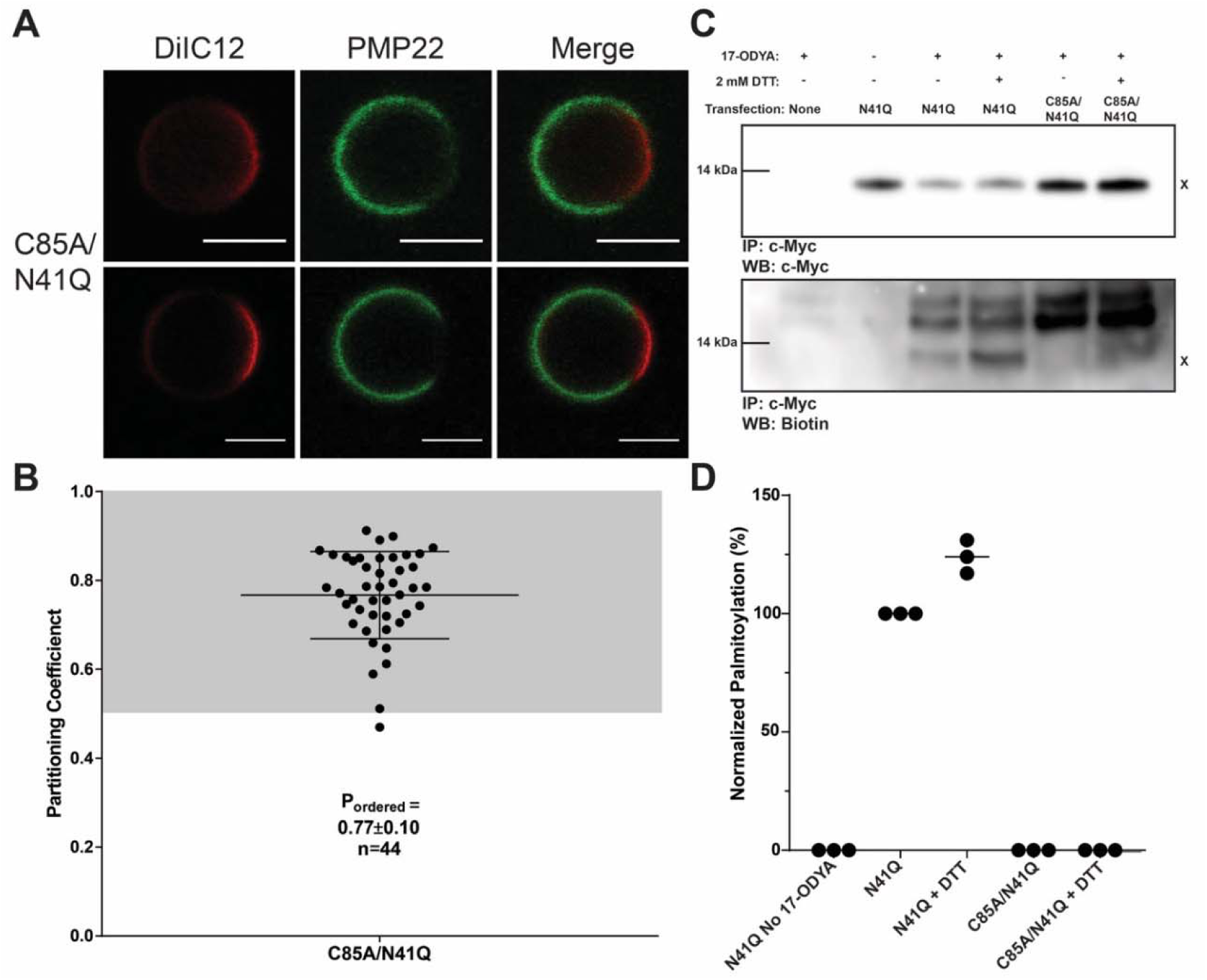
Palmitoylation is not required for PMP22 localization to the ordered phase. **(A)** Upper and lower triple images are representative examples of PMP22-containing GPMVs from C85A/N41Q PMP22-transfected HeLa cells. Scale bar, 5 µm. **(B)** Quantification of C85A PMP22 partitioning coefficients from three independent biological experiments. Mean ± SD is reported and plotted on the graph. **(C)** Representative western blots of PMP22 and palmitoylated PMP22 from cells treated with or without 2 mM DTT. The top blot shows PMP22 immunopurified from cell lysates while the bottom blot shows palmitoylated protein from the immunopurified lysates which was identified via addition of a biotin handle to alkyne palmitate. **X** marks the PMP22 signal. Uncut western blots shown in **Figure S5**. **(D)** Quantification of the amount of palmitoylated PMP22 from three biological replicates. The amount of palmitoylated PMP22 from each sample is quantified by dividing the intensity from the biotin blot by the intensity of the myc blot and then normalized to the amount of palmitoylated PMP22 in the N41Q sample without DTT treatment.

### Cholesterol binding motifs in PMP22 do not mediate its ordered phase preference

We next tested whether either or both of the predicted cholesterol binding sites in PMP22 play a role in its ordered phase preference. It has been shown that *pmp22 -/-* Schwann cells exhibit reduced plasma membrane levels and abnormal localization of cholesterol (*30, 32*). These cells also show reduced migration, adhesion, and lamellipodia extension, all of which can be restored through external supplementation of cholesterol in the culture media. PMP22 contains both a classical cholesterol-recognition amino acid consensus (CRAC) motif in TM4 (L-X_1-5_-Y-X_1-5_-K) and an inverted CRAC (CARC) motif in TM3 (K-X_1-4_-Y-X_1-6_-I), as illustrated in orange in **Fig. 1A**. While these motifs are loosely defined and are not always indicative of direct cholesterol interaction(*45*), there is substantial experimental and computational evidence supporting the notion that these motifs are sometimes directly involved in binding cholesterol (*46, 47*).

We mutated one or both of the essential Tyr residues in the CARC and CRAC motifs to Ala (Y97A, Y153A, and Y97A/Y153A mutants; **Fig. 1A,** purple). We then assessed the phase preference for each mutant in GPMVs. Mutation of Tyr97 had no effect on PM levels of PMP22 but mutation of Tyr153 or of both Tyr residues led to decreased PMP22 levels at the PM, suggesting lower expression and/or surface trafficking efficiency for these mutants (**Fig. S2).** Nevertheless, the mutations did not significantly alter the ordered phase preference of PMP22 (**Fig. 3A-B)**. Y97A exhibited a P_ordered_ of 0.83±0.10, Y153A exhibited a P_ordered_ of 0.81±0.13, and the double mutant Y97A/Y153A yielded a P_ordered_ of 0.71±0.15. P_ordered_ for the double mutant is only slightly reduced. We interpret these results to indicate that the presence of CRAC and/or CARC motifs are not significant drivers of the preference of PMP22 for ordered phase domains.

**Figure 3.**
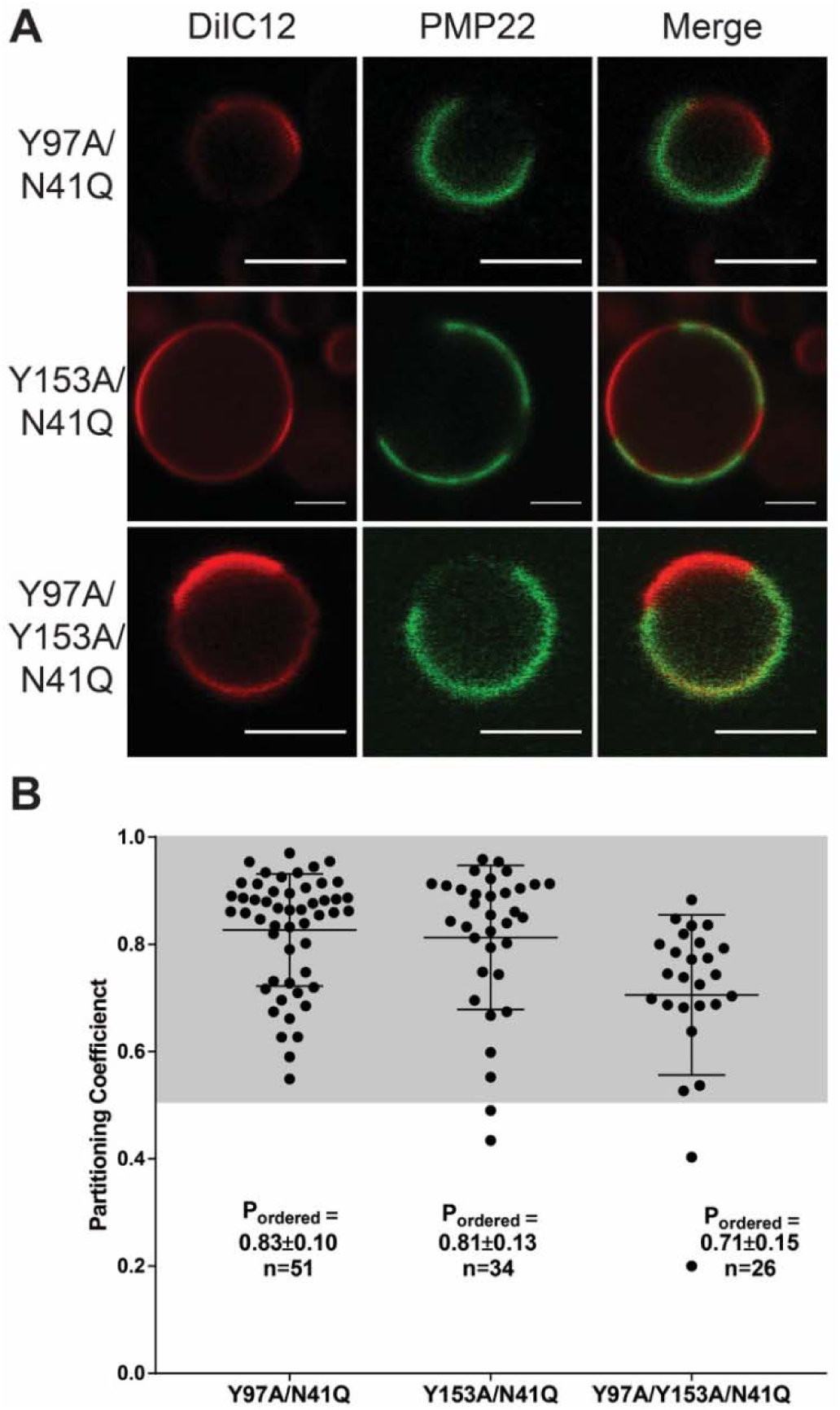
Cholesterol interaction motifs do not contribute to the ordered phase domain preference of PMP22. **(A)** Representative PMP22 containing GPMVs from Y97A/N41Q, Y153A/N41Q, and Y97A/Y153A/N41Q mutant forms of PMP22-transfected HeLa cells. Scale bar, 5 µm. **(B)** Quantification of PMP22 partitioning coefficients from three independent biological experiments. Mean ± SD is reported and plotted on the graph.

### Possible role for TM1 in determining the phase preference of PMP22

Since palmitoylation and cholesterol interaction motifs do not appear to play a major role in defining the phase preference of PMP22, we hypothesized that there is something intrinsic to the structure of the protein that drives its preferential association with the ordered phase. Perhaps the most unusual structural feature of PMP22 is its first transmembrane helix (TM1), which is relatively long at 26 residues and lacks any polar residues at its N-terminus. The first 6 residues of PMP22 are also the first 6 residues of TM1: Met-Leu-Leu-Leu-Leu-Leu-. It is odd for the absolute N-terminus of a membrane protein to be part of the first TM segment.

We conducted genetic “intolerance analysis” of human PMP22 to measure the missense tolerance ratios (MTR), (*48, 49*) for all possible segments of PMP22. MTR vs. sequence plots report how tolerant a region of a protein is to the possible evolutionary purifying effects of missense mutations, as judged based on analyzing all sequences for that gene in the >200,000 currently available human exome or whole genome sequences. High MTR values point out which regions of the protein are tolerant to missense mutations. Low MTR values indicate regions of the protein for which genetic variations leading to amino acid changes are rarely observed in the population, meaning that the DNA is *intolerant* to missense mutations. **Fig. 4A** shows MTR analysis results for PMP22.

**Figure 4.**
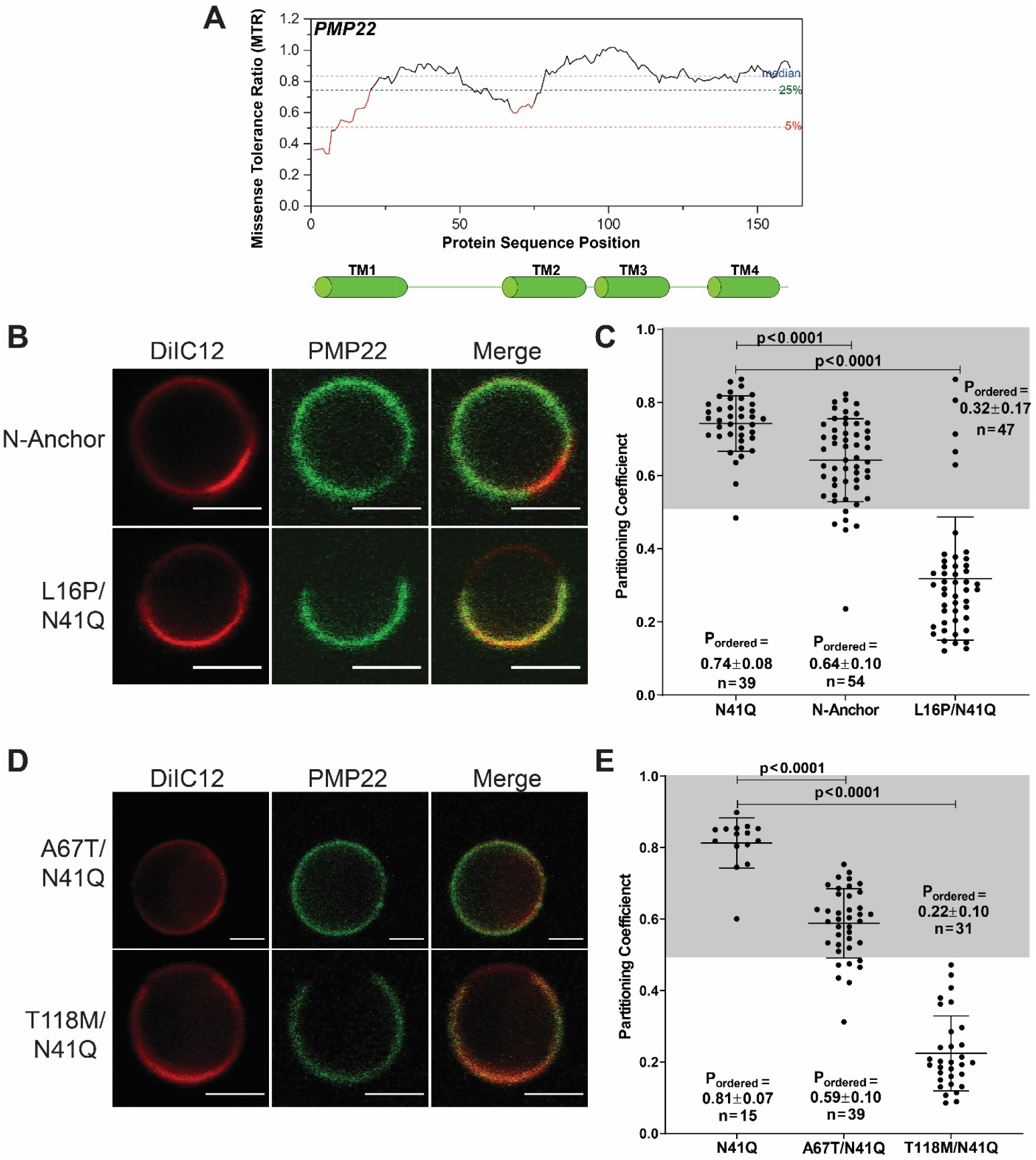
TM1 plays a role in conferring ordered phase domain preference to PMP22. **(A)** Missense tolerance analysis of *pmp22.* The MTR for a 31 residue sliding window of *pmp22* is shown via the black and red solid line plot. MTR values of 1.0 indicate that the frequency of missense mutations encoding an amino acid change in that segment is the same as seen for silent missense mutations in the same segment. An MTR value lower than 1.0 indicates the segment is subject to “purifying selection” meaning some degree of evolutionary intolerance within that segment to amino acid-changing mutations. MTR values highlighted in red are those for which the false discovery rate (FDR) is less than 0.1, indicating the deviation of the MTR value from 1 is statistically robust(*48–50*). **(B)** Representative PMP22-containing GPMVs from N-anchor mutant PMP22 and L16P/N41Q PMP22-transfected HeLa cells. Scale bar, 5 µm. **(C)** Quantification of the PMP22 partitioning coefficients of N41Q, N-Anchor, and L16P/N41Q mutants from three paired biological experiments. **(D)** Representative PMP22-containing GPMVs from A67T/N41Q and T118M/N41Q PMP22-transfected HeLa cells. **(E)** Quantification of the PMP22 partitioning coefficients of N41Q, A67T/N41Q, and T118M/N41Q mutants from three paired biological experiments. Mean ± SD is reported and plotted on the graph. The non-parametric Mann-Whitney U test was used for all statistical analysis

Overall, PMP22 is seen to be generally tolerant of missense mutations, reflecting the fact that while mutations in PMP22 cause disease, the disease is not fatal and usually does not eliminate reproductive capacity. However, the one region of the protein seen to be relatively intolerant to gene variations in the human population is the N-terminus of TM1 (red trace **Fig. 4A**). The combined importance and distinctive features of TM1 led to the hypothesis that this segment might be important in helping to define the ordered phase domain preference of PMP22. We first tested the effect of adding a conventional cytosolic anchor to the TM1 segment of PMP22. A positively charged segment (-Arg-Arg-Gly-Gly-) was added between the native N-terminal Met residue and the penta-Leu repeat follows at the start of TM1. This generated an “N-Anchor” mutant form of PMP22 (**Fig. 1A**, N-Anchor). The addition of this anchor was seen to have no effect on PMP22 cellular localization, as this variant predominately localized to the cell surface in fixed cell images similar to WT-like N41Q PMP22 (**Fig. S2**). When we then measured the phase preference of the N-Anchor mutant in GPMVs we observed a modest but significant reduction in P_ordered_ to 0.64±0.11 (**Fig. 4B-C**) compared to N41Q PMP22 (P_ordered_=0.74±0.08 in experiments performed in parallel). Addition of the N-anchor to PMP22 reduces but does not eliminate its preference for the ordered phase.

We next tested the phase preference of PMP22 containing a disease mutation in TM1: the L16P variant (**Fig. 1A**, red), which is also known as the “Trembler-J” mutation because of its mouse phenotype(*28, 29, 41*). Under WT/mutant heterozygous conditions L16P PMP22 causes severe demyelination in both human and mice. Moreover, biophysical studies on L16P PMP22 in detergent micelles showed that the L16P mutation introduces a flexible hinge in the helix, destabilizing the fully folded form of the protein and causing it to adopt an unfolded or folding intermediate state in which TM1 is disassociated from TM2-4, which remain bundled, but as a molten-globule (*41, 50, 51*). While introduction of the L16P mutation exhibits significantly reduced cell surface expression compared to N41Q PMP22 (**Fig. S2**), we still were able to generate enough L16P PMP22 containing GPMVs to measure its phase preference. For L16P PMP22 we determined P_ordered_ to be 0.32±0.17 (**Fig. 4B-C**). Thus, the L16P mutation dramatically reverses the phase preference of PMP22 such that the protein now prefers to partition into disordered membrane phase domains. This result suggests that the formation of stable tertiary structure in WT PMP22 is important for its ordered phase preference.

Since the CMTD causing L16P mutation was shown to reverse the membrane phase preference of PMP22, we decided to test the phase partitioning of two other disease-causing mutations located in other transmembrane segments. The A67T point mutation is located at a site near the exoplasmic end of TM2. This mutation is known to induce a slight destabilization of the conformational stability of PMP22 (*41*) and causes a mild form of CMTD, hereditary neuropathy with liability to pressure palsies (HNPP) (*28*). In GPMVs A67T PMP22 exhibited a P_ordered_ of 0.59±0.10, a significant reduction of ordered phase preference compared to PMP22 (0.81±0.07, as measured in parallel experiments; **Fig. 4D-E**). The T118M point mutation destabilizes the conformational stability of PMP22 to a much greater extent compared to A67T (*41*), but less so than the L16P mutation, and is known to cause a moderate form of CMTD (*28*). In GPMVs, T118M PMP22 displayed a P_ordered_ of 0.22±0.11, reflecting pronounced preference for disordered membrane phases (**Fig. 4D-E**). These results suggest that the tertiary structural stability of PMP22 is essential to the affinity of WT PMP22 for ordered membrane domains— merely inserting all TM segments of the protein into the membrane correctly is not enough.

### PMP22 alters the biophysical properties of GPMVs and promotes formation of ordered phase domains

Because it has been shown that PMP22 is critical for the formation of stable membrane domains in Schwann cells (*30*), we hypothesized that PMP22 may alter the stability of phase separation between ordered and disordered phase domains in GPMVs. To test this, we first determined the miscibility temperature (T_Misc_) of PMP22-containing GPMVs compared to GPMVs derived from cells transfected with an empty vector (“MOCK” conditions; **Fig. 5A**). T_Misc_ is defined as the temperature at which 50% of the GPMVs exhibit phase separation (*9, 12, 52*). A higher T_Misc_ suggests more stable phase-separated membrane domains. We collected images of >100 GPMVs at each temperature over temperatures ranging from 12.5°C to 32.5°C and calculated the fraction of phase-separated GPMVs at each temperature. Fitting this data to a sigmoidal curve allowed us to determine the T_Misc_ of the GPMVs. **Fig. 5B** shows the T_Misc_ seen for PMP22-containing GPMVs and for GPMVs from MOCK transfected cells. PMP22-containing GPMVs exhibited a T_Misc_ of 20.2±0.6 °C whereas MOCK GPMVs showed a T_Misc_ of 18.8±0.4°C. The T_Misc_ seen for empty GPMVs was similar to what has previously been reported (*9, 52*). To validate that the increase in T_Misc_ was not due to general overexpression of a TM protein at the plasma membrane, we measured the T_Misc_ of tgLAT containing GPMVs (**Fig. S6**). GPMVs containing tgLAT exhibited a T_Misc_ of 18.4±0.4 °C, similar to MOCK conditions (**Fig. 5B**), suggesting that the T_Misc_ increase of PMP22-containing GPMVs was not due to generic membrane protein overexpression. This result indicates that PMP22 stabilizes phase separation in GPMVs.

**Figure 5.**
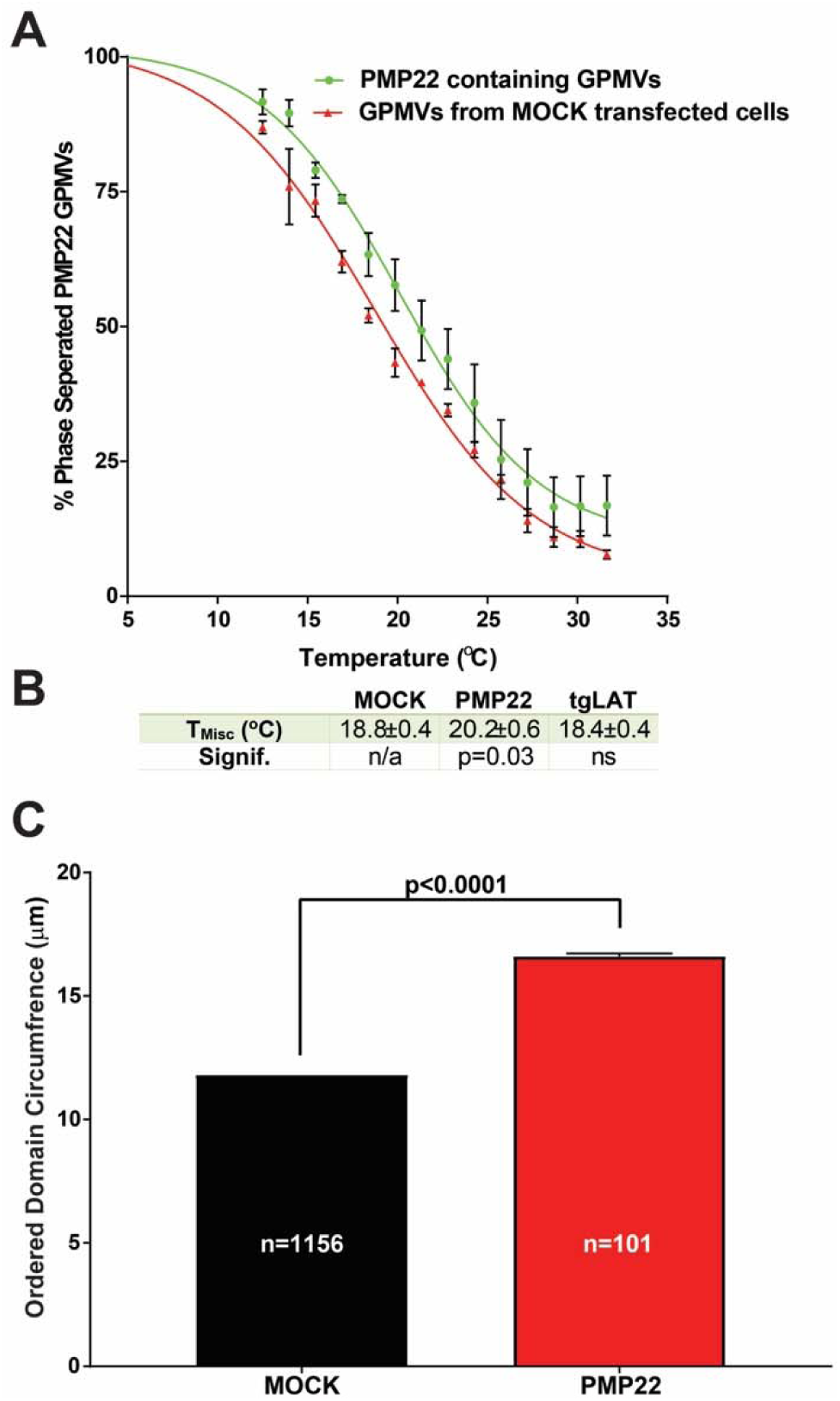
PMP22 alters the biophysical properties of GPMVs. **(A)** Percent phase separation of GPMVs containing N41Q PMP22 (green) or of GPMVs derived from cells transfected with an empty vector (MOCK; red) at various temperatures. Each point shows the average of three independent experiments and the error bars represent the standard error of the mean (SEM). >100 GPMVs were measured at each temperature in each experiment. Plots are fit to a sigmoidal curve. **(B)** Calculation of the phase miscibility temperature (T_Misc_) for GPMVs from MOCK transfected cells, GPMVs containing PMP22, and GPMVs containing tgLAT. T_Misc_ is calculated from the fit of the sigmoidal curve for each independent experiment. The reported value represents the mean T_Misc_ ± SEM. Significance was determined using a non-parametric Mann-Whitney U test comparing PMP22-continaing GPMVs or tgLAT-containing GPMVs to MOCK GPMVs. **(C)** Ordered domain sizes (μm) from GPMVs obtained from cells transfected with either an empty vector (MOCK) or GPMVs containing N41Q PMP22. Data was obtained for three biological replicates for MOCK samples and six biological replicates for PMP22. Significance was determined using a non-parametric Mann-Whitney U test.

We also examined whether the presence of PMP22 increased the size of ordered phase domains in GPMVs. To do this we imaged a large number of GPMVs derived from cells transfected with either an empty vector or one encoding PMP22. For these images we measured the radii of individual GPMVs as well as the fraction of each GPMV that contained the disordered phase maker DiIC12. From this information we were able to calculate the relative size of ordered domains in GPMVs, as indicated in **Fig. 5C** (see **Fig. S7** for raw values). We found that GPMVs derived from cells transfected with an empty vector contained ordered domains with an average circumference of 11.8±0.1 μm whereas GPMVs with PMP22 contained ordered domains with a circumference of 16.6±0.1 μm. This significant increase in size suggests that PMP22 is able to stabilize the ordered membrane domains of GPMVs. Combined with the fact that PMP22 increases the T_Misc_ of GPMVs, we conclude that PMP22 is able to alter the biophysical properties of GPMVs to promote formation of ordered phase domains.

## Discussion

While there are a number of biochemical reports that identify membrane proteins founds to co-localize with biochemically isolated “lipid raft” cell fractions (*53–55*), to the best of our knowledge this is the first work that *quantitively* demonstrates the ordered phase membrane domain preference of a multi-pass membrane protein in cell-derived GPMVs.

### Factors contributing to the ordered phase domain preference of PMP22

Much effort has been devoted to understanding the driving forces of membrane phase preference for single-pass transmembrane proteins. Previous work culminated in an elegant paper published by the Levental group that provided a quantitative model describing the biochemical and biophysical features promoting ordered phase domain partitioning for single-pass transmembrane proteins (*20*). Ordered phase domain partitioning of single-pass proteins is promoted by increasing length for the transmembrane helix, by the presence of one or more palmitoyl chains, and by the presence of small amino acid side chains in the fully membrane-exposed TM segment, especially for the half that sits in the exoplasmic bilayer leaflet. Whether components of this model can be extrapolated to ordered phase-preferring multi-pass membrane proteins is not yet clear. However, this study of PMP22 represents an important step in exploring this question.

The results demonstrated that the tetraspan integral membrane protein PMP22 has a distinct preference to partition into ordered phase membrane domains of GPMVs derived from both HeLa and primary Schwann cells. Unlike ordered phase-preferring single-pass transmembrane proteins (*20, 23, 24*), partitioning of PMP22 is not driven mainly by palmitoylation. Even though native PMP22 is palmitoylated, it retains its strong preference for the ordered phase even under conditions in which its palmitoylation site is mutated away. Our results demonstrate that this modification is not required for the ordered membrane phase preference of PMP22, as it seems to be for single-pass membrane proteins. Ordered phase partitioning of PMP22 was also found not to be associated with the putative CARC and CRAC cholesterol binding motifs present in PMP22.

The PMP22 TM1 segment appears to play a specific role in determining its phase preference, as addition of a hydrophilic anchor to the apolar N-terminal end of this helix (which also serves as the N-terminus for the entire protein) significantly reduced the preference of PMP22 for the ordered phase. We speculate that the lack of polar residues flanking the start of TM1 may enable WT PMP22’s N-terminus to adapt more easily to the more rigid matrix and thicker transbilayer span of ordered phase membranes relative to the adjacent disordered phase. Future work will be needed to test this possibility and explore possible alternative explanations. We also observed that introduction of the L16P mutation in the middle of TM1 reversed the phase preference of PMP22 so that it now favors the disordered phase. We have previously shown that the L16P mutant converts the straight and uninterrupted WT TM1 helical segment into a pair of helices linked by a flexible hinge (*50, 51*). This causes TM1 to dissociate from the other TM helices to favor a destabilized form of the protein in which TM2, TM3, and TM4 remain in contact as a molten globular bundle while TM1 is dissociated in the membrane, tethered to the rest of the protein by the TM1-TM2 loop.

In addition to L16P PMP22, we also tested the phase preference of other CMTD PMP22 mutant forms, A67T and T118M. We previously observed that each of these mutant forms of PMP22 are destabilized relative to WT (*41*). Interestingly, we found that the A67T mutation, which induces only a modest decrease of the conformational stability of PMP22, also caused only a modest decrease in its preference for the ordered phase. A more severely destabilizing mutation, T118M, resembled L16P (also very destabilizing) by dramatically shifting its phase preference to favor the disordered membrane phase. For these mutants we hypothesize that this shift in phase preference is the consequence of the global destabilization of the tertiary fold of the protein, even though the unfolded form retains its correct tetraspan membrane topology.

That formation of correct tertiary structure by PMP22 is required for ordered phase partitioning indicates that that the folded structure of PMP22 has traits that favor ordered phase partitioning relative to the disordered phase. Examination of a homology/Rosetta model for the structure of PMP22 (*50*) suggest that two of its transmembrane helices may be longer than average (at least 26 residues each) and that the transmembrane domain has a fairly featureless surface. While there does not seem to the general preponderance of residues with small side chains in the exoplasmic half of the PMP22 transmembrane domain as appears to be a feature of ordered phase-preferring single-pass membrane proteins (*20, 56*), the presence of a Ser-Ala-Ala-Ala segment at the exoplasmic end of TM3 is intriguing. The intracellular “domain” of PMP22 is another distinctive feature, being comprised only of the N-terminal amino group of Met1, a 4 residue loop connecting TM2 and TM3, and four charged residues that follow TM4. Testing whether any of these features are contributing factors to PMP22’s ordered phase preference will require many additional experiments, which we hope are motivated by the result of this paper.

### The preference of PMP22 for ordered phase membrane domains of cell-derived GPMVs does not extend to the L_o_ phase of synthetic lipid vesicles

In previous work we showed that purified recombinant PMP22 can be reconstituted into giant unilamellar vesicles (GUVs) containing synthetic lipids (*57*). Under GUV conditions in which the synthetic lipids separated into ideal L_d_ and L_o_ lipid phases, it was observed that PMP22 partitioned exclusively to the disordered L_d_ phase. Why are the results for PMP22 partitioning in GPMVs at odds with what was observed in GUVs? Our results show that neither of the known post-translational modifications of PMP22, N-glycosylation at N41 or S-palmitoylation at C85, are required for the ordered phase preference of PMP22 in GPMVs, such that it can be ruled out that the lack of post-translational modifications of the recombinant PMP22 used in the earlier GUV studies is the basis for its L_d_ phase preference. We suggest instead that the variance between the results from the GPMV and GUV studies point to the fact that the difference in order between phase-separated domains in cell-derived GPMVs is much reduced relative to GUVs comprised of a well-defined ternary mixture of synthetic lipids. This phenomenon has previously been documented for single span membrane proteins (*8, 18, 58*). Given that the P_ordered_ for PMP22 in GPMVs was seen in this work to be nearly 0.8 means that the energy by which PMP22 favors the ordered phase over the disordered phase in GPMVs is on the order of –RTln(4) = −0.8 kCal/mol. One can easily imagine that the highly-ordered packing that occurs in ideal L_o_ phases (but only to a much lesser degree in GPMVs) would need to be disrupted to accommodate partitioning of a membrane protein and that this unfavorable energy contribution could easily reverse the overall energetics of partitioning in GUVs to favor the L_d_ phase.

### PMP22 stabilizes ordered phase domains and promotes their formation

While some proteins are thought to passively associate with raft-like domains, others can actively promote their formation by clustering raft components and stabilizing ordered domains(*25, 59*). Proteins that modulate membrane order or fluidity also are capable of regulating phase separation(*60, 61*). Here, we identify PMP22 as a new example of a protein that can directly stabilize ordered phase membrane domains. PMP22-containing GPMVs exhibited a higher T_Misc_ than GPMVs containing tgLAT or cells transfected with an empty vector (**Fig. 5A-B, Fig. S5**). That phase separation persists at higher temperatures in GPMVs containing PMP22 suggests that this protein can directly stabilize ordered phase membrane domains (*9*). This is consistent with previous results from studies of *pmp22 -/-* mice showing that the distribution of molecules typically associated with ordered phase membrane domains (such as cholesterol, and GM1 ganglioside) are decreased at the plasma membrane (*30*). Moreover, Schwann cells isolated from these mice showed elongation and migration defects that could be corrected by external supplementation of the culture medium with cholesterol. Additionally, our results suggest that PMP22 is able to promote ordered domain formation. We showed that GPMVs containing PMP22 had ordered membrane domains on average ∼5 μm larger than those in GPMVs without PMP22 (**Fig. 5C**). This may be due to an increased concentration in cholesterol in PMP22-containing GPMVs since it has recently been shown that PMP22 is regulates cholesterol PM trafficking (*32*). It seems likely that the mechanisms underpinning this regulatory function of PMP22, as well as its ability to promote ordered phase formation, is closely related to its preference to partition into ordered membrane phase domains.

## Conclusions

We have documented PMP22 as the first multi-pass membrane protein to exhibit a preference to partition into the ordered phase of cell-derived GPMVs. This phase preference appears to be closely linked to its unusual TM1 and also requires not just correct membrane integration of its four transmembrane helices, but formation of correct tertiary structure. Additional experiments will be required to determine exactly what features of its folded structure confer its preference for the ordered phase. Moreover, it remains unclear just how many other multi-pass membrane proteins will share the phase domain preference of PMP22 and whether they will resemble PMP22 in terms of driving traits. It is hoped that the results of this work will inspire future studies of both PMP22 and other multi-pass membrane proteins to address these issues.

## Materials and Methods

### Materials

DiIC12 was purchased from Life Technology (Eugene, USA). NBD-PE was purchased from Avanti (Alabaster, USA). Anti-myc AlexaFluora-647, anti-biotin, and anti-myc were purchased from Cell Signaling Technologies (Danvers, USA). DTT, tris(2-carboxyethyl)phosphine (TCEP), and 4-(2-hydroxyethyl)-1-piperazineethanesulfonic acid (HEPES) were purchased from Research Products International (Mount Prospect, USA); DTT was prepared fresh for every use. Paraformaldehyde (PFA) was purchased as a 16% stock solution from Electron Microscopy Sciences (Hatfield, USA). CaCl_2_, NaCl, CuCl_2_, Tris[(1-benzyl-1H-1,2,3-triazol-4-yl)methyl]amine (TBTA), anti-myc magnetic beads and propidium iodide were purchased from Fisher Scientific (Fair Lawn, USA). 17-Octadecynoic Acid (17-ODYA) and biotin-azide were purchased from Cayman Chemicals (Ann Arbor, USA). HeLa cell lines were acquired from ATCC (Manassas, USA). Primary rat Schwann cells (RSCs) were a generous gift from the lab of Dr. Bruce Carter at Vanderbilt University.

### Cloning

Human cDNA for PMP22 was subcloned into a pCDNA3.1 mammalian expression vector. To make PMP22 immunologically detectable we used QuickChange mutagenesis to insert a myc epitope into the second extracellular loop of PMP22 within the pCDNA3.1 vector(*41*). QuickChange mutagenesis was also used to introduce point mutations and insert the ‘N-Anchor’ sequence (-RRGG-) between Met1 and Leu2. Plasmids were purified using a GenElute HP Plasmid MidiPrep Kit (Sigma-Aldrich). The tgLAT construct(*20*) used in these studies was a generous gift from the lab of Dr. Ilya Levental at the McGovern Medical School, University of Texas.

### Cell Culture and Transfections

HeLa and RSCs were cultured in Dulbecco’s modified Eagle medium (DMEM) containing 10% fetal bovine serum (FBS) and 1% pen/strep at 37°C and 5% CO_2_. Culture medium for RSCs was supplemented with 2 µM forskolin (Sigma-Aldrich). ∼24 hours prior to transfection, cells were plated so as to be 40-50% confluent at the time of transfection. Cells were transfected using FuGene Transfection Reagent (Promega, Madison USA) with a FuGene:DNA ratio of 3:1 in OptiMEM. 6 cm^2^ plates were transfected with 1.5 µg DNA. The transfection medium was removed from cells ∼12-15 hours post-transfection and cells were washed with DPBS and fresh culture media was added to each plate.

### GPMV Preparation

36 hours after transfection, the medium was removed from cells and cells were washed three times with inactive GPMV buffer (10 mM HEPES, 150 mM NaCl, 2 mM CaCl_2_ pH 7.4). Cells were consistently 70-80% confluent at the time of GPMV prep. Active GPMV buffer (GPMV buffer plus 2 mM DTT and 25 mM formaldehyde) was then added to the plates, and cells were incubated at 37° C with gentle shaking (70 RPM) for 90 minutes. DiIC12 or NBD-PE was then added to the plates from a stock solution of 0.5 mg/mL in EtOH to a final concentration of 0.5 µg/mL and cells were gently rocked at room temperature for 15 minutes. The GPMV-containing supernatant was then decanted into 1.5 mL Eppendorf tubes, and an anti-myc AF647 mAb was added to the solution (1:750 uL dilution) and gently agitated in the dark at room temperature for at least 3 hours. GPMVs were then allowed to settle in the dark to the bottom of the tube at 4°C for 2-24 hours (we observed no difference in GPMV quality whether we imaged immediately or at 24-hour post GPMV prep). 30 minutes prior to imaging, 270 µL of GPMV solution was pipetted from the bottom of the Eppendorf tube and sandwiched between 2 coverslips coated with 0.1% BSA and separated by a 0.5 mm thick silicone isolator (Electron Microscopy Sciences).

### GPMV Imaging

GPMVs were imaged using a Zeiss LSM 510 confocal microscope using a 1.2 NA Zeiss Plan-Neofluor 40X objective. The confocal pinhole was set to 150 nm for all experiments. The fluorophores were excited using the 488 nm line of a 40 mW argon laser (NBD-PE, mEGFP, and AF-488), the 543 nm line of a HeNe laser (DiIC-12, propidium iodide), or the 633 nm line of a HeNe laser (AF-647). Images were collected at a 1X digital zoom for the case of miscibility temperature measurements and at 8-10X digital zoom for quantifying phase partitioning with a 512×512 pixel resolution. The stage was cooled using a Linkan Peltier Cooling system (Tadworth, UK).

### Quantifying GPMV phase partitioning

Fig. S1 shows a representative example of how P_ordered_ was calculated. Briefly, GPMVs were labeled with either a disordered membrane phase marker (DiIC-12) or an ordered membrane phase marker (NBD-PE). PMP22-containing GPMVs were then labeled with anti-myc AF647-labeled antibodies. To determine the phase partitioning of PMP22, GPMVs were imaged in the green or red (NBD-PE or DiIC-12 respectively) and far-red channels, sequentially. To determine the phase partitioning of tg-LAT, GPMVs were imaged in the green and red channels sequentially. Line scans across a single GPMV were performed in all channels using the ImageJ software to determine the fluorescent intensity at every pixel. The position of the line was set so that it intersected with both an ordered and disordered region of the GPMV using the DiIC12 or NBD-PE channels as the references. This same line was used to measure the intensity in the protein (PMP22 or tgLAT) channel. The line scans were smoothened using a moving average (10 pixels) in Microsoft Excel. A partitioning coefficient P_ordered_ was then calculated as previously described(*25*) as

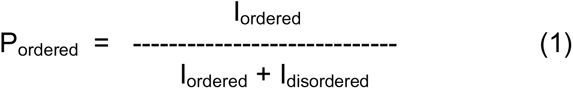

where I_ordered_ and I_disordered_ are the fluorescence intensity of the protein channels in the ordered and disordered phases, respectively. Three independent lines were chosen for each GPMV and the mean P_ordered_ was calculated and reported for individual GPMVs.

### Quantification of Palmitoylation using ‘Click’ chemistry

∼24 hours after cells were transfected, cells were incubated overnight in media containing 100μM 17-ODYA and a 1% final concentration of DMSO (or just DMSO for no 17-ODYA control). Cells were then incubated for 90 minutes with or without 2 mM DTT. Following incubation, cells were lysed for 1 hour at 4°C in 150 μL lysis buffer (50 mM Tris, 150 mM NaCl_2_, 0.3% CHAPS, 0.1% SDS, pH 7.4). Lysates were cleared via centrifugation for 15 mins at 14,000xg. Protein concentrations were determined via Bradford assay and 75 μg total protein was added to 10 μL of anti-myc conjugated magnetic beads for each lysate. Volumes for each lysate were brought up to 150 μL total in lysis buffer and beads and lysates were incubated with end-over-end rotation at 4°C overnight. The following day, beads were washed three times with lysis buffer and bound proteins were eluted with 25 μL of elution buffer (50 mM HEPES, 150 mM NaCl, 2% SDS pH 7.0) and transferred to fresh tubes. The following was added to the eluents: biotin azide to a final concentration of 20 μM, TCEP to a final concentration of 1 μM, 20 μM TBTA, and 2 μM CuSO_4_. Reactions were mixed at room temperature with end-over-end rotation for 2 hours before being quenched via the addition of EDTA to a final concentration of 1 mM. Samples were then split in two and analyzed via western blotting for PMP22 (1:8000 dilution of c-myc antibody) and biotin (1:1000 dilution of biotin antibody).

### Measurement of Miscibility Temperature (T_Misc_)

5×5 tile scans of GPMV samples were imaged at 1X digital zoom at temperatures ranging from 12.5° C to 32.5°C. Images were then randomized and GPMVs were blindly and manually classified as being either phase-separated or containing a single uniform phase. The fraction of vesicles that were phase separated at each temperature was then calculated. Plotting %-phase separated versus temperature yielded a curve that was fit to a sigmoidal function and the miscibility temperature (T_Misc_) was defined as the temperature at which 50% of GPMVs were phase separated. Three independent biological experiments were performed and >100 GPMVs were imaged and classified for each temperature of each repeat. Classifications were performed blindly to the temperature at which the images were collected.

### Fixed-Cell Imaging

Cells were cultured on glass coverslips and transfected as described above. 36 hours post transfection, cells were fixed with 4% PFA and permeabilized with 0.25% Triton X-100. Cells were treated with 1 ug/mL of RNase (Sigma-Aldrich) and PMP22, as then detected with mouse anti-myc (1:500) and visualized with donkey anti-mouse AF-488 (Invitrogen; 1:500). Nuclei were stained with propidium iodide (Sigma-Aldrich; 0.33 µg/mL). Coverslips were mounted to a slide using ProLong Gold (Invitrogen) and allowed to dry for 24 hours. Cells were then imaged using the 40X objective on the Zeiss LSM 510 confocal microscope at an optical zoom of 3-4X. The optical slice was set to <1µM in each image.

### Calculation of GPMV Ordered Domain Size

Images of GPMVs derived from cells transfected either with an empty or N41Q PMP22 pCDNA3.1 vector were collected in a high throughput manner on an ImageXpress Micro XL (Molecular Devices) using a 40X objective. MATLAB (MathWorks) was used to calculate the radius of each GPMV in pixels, and the percentage of GPMVs in the ordered phase using the DiIC12 dye to mark the ordered phase. The radius was converted from pixel to μm using the conversion factor of 0.34 μm:1 pixel for the 40X objective. The circumference of each GPMV was then calculated using the equation: circumference=2π*radius, and ordered domain size was calculated by multiplying the fraction of each GPMV that was in the ordered phase by its circumference.

### Statistical Analysis

The nonparametric Mann-Whitney U test was used to compare pairs of data. Significance is reported when determined. GraphPad Prism was used to perform all statistical analysis.

### Missense Tolerance Ratio

The MTR plot for PMP22 was generated using the online software MTR Viewer Version 2 (http://biosig.unimelb.edu.au/mtr-viewer/).

## Acknowledgments

We thank Dr. Krisnan Raghunathan and Dr. Ajit Tiwari for introduction to the GPMV technique, Arina Hadziselimovic for assistance with cloning, the Vanderbilt Cell Imaging Shared Resource (CISR) for access to the confocal microscope, the Vanderbilt Institute of Chemical Biology and High Throughput Screening (HTS) facility, and Dr. Hui Huang for access and assistance to the ImageExpress, the lab of Bruce Carter at Vanderbilt for harvesting and help culturing RSCs, and Dr. Melani Ohi (University of Michigan) for critical early revisions to the manuscript.

## Funding

This work was supported by US NIH grant R01 NS095989 and a Vanderbilt Stanley Cohen Award, both awarded to both CRS and AKK. JTM was supported by NIH fellowship F31 NS113494 and NIH training grant T32 NS00749. The CISR is supported by NIH grants CA68485, DK20593, DK58404, DK59637, and EY08126.

## References

1. K. Simons, E. Ikonen, Functional rafts in cell membranes. Nature 387, 569–572 (1997).

2. D. A. Brown, E. London, Functions of lipid rafts in biological membranes. Annu Rev Cell Dev Biol 14, 111–136 (1998).

3. M. L. Kraft, Plasma membrane organization and function: moving past lipid rafts. Mol Biol Cell 24, 2765–2768 (2013).

4. S. L. Veatch, S. L. Keller, Separation of liquid phases in giant vesicles of ternary mixtures of phospholipids and cholesterol. Biophys J 85, 3074–3083 (2003).

5. D. G. Ackerman, G. W. Feigenson, Lipid bilayers: clusters, domains and phases. Essays Biochem 57, 33–42 (2015).

6. X. Cheng, J. C. Smith, Biological Membrane Organization and Cellular Signaling. Chem Rev, (2019).

7. E. Sezgin, I. Levental, S. Mayor, C. Eggeling, The mystery of membrane organization: composition, regulation and roles of lipid rafts. Nat Rev Mol Cell Biol 18, 361–374 (2017).

8. I. Levental, S. Veatch, The Continuing Mystery of Lipid Rafts. J Mol Biol 428, 4749–4764 (2016).

9. S. L. Veatch et al., Critical fluctuations in plasma membrane vesicles. ACS Chem Biol 3, 287–293 (2008).

10. A. R. Honerkamp-Smith, S. L. Veatch, S. L. Keller, An introduction to critical points for biophysicists; observations of compositional heterogeneity in lipid membranes. Biochim Biophys Acta 1788, 53–63 (2009).

11. E. Gielen et al., Rafts in oligodendrocytes: evidence and structure-function relationship. Glia 54, 499–512 (2006).

12. T. Baumgart et al., Large-scale fluid/fluid phase separation of proteins and lipids in giant plasma membrane vesicles. Proc Natl Acad Sci U S A 104, 3165–3170 (2007).

13. M. Aureli, S. Grassi, S. Prioni, S. Sonnino, A. Prinetti, Lipid membrane domains in the brain. Biochim Biophys Acta 1851, 1006–1016 (2015).

14. D. A. Brown, Preparation of detergent-resistant membranes (DRMs) from cultured mammalian cells. Methods Mol Biol 1232, 55–64 (2015).

15. D. A. Brown, Lipid rafts, detergent-resistant membranes, and raft targeting signals. Physiology (Bethesda) 21, 430–439 (2006).

16. H. Heerklotz, Triton promotes domain formation in lipid raft mixtures. Biophys J 83, 2693–2701 (2002).

17. S. Schuck, M. Honsho, K. Ekroos, A. Shevchenko, K. Simons, Resistance of cell membranes to different detergents. Proc Natl Acad Sci U S A 100, 5795–5800 (2003).

18. K. R. Levental, I. Levental, Giant plasma membrane vesicles: models for understanding membrane organization. Curr Top Membr 75, 25–57 (2015).

19. J. H. Lorent, I. Levental, Structural determinants of protein partitioning into ordered membrane domains and lipid rafts. Chem Phys Lipids 192, 23–32 (2015).

20. J. H. Lorent et al., Structural determinants and functional consequences of protein affinity for membrane rafts. Nat Commun 8, 1219 (2017).

21. E. Sezgin et al., Elucidating membrane structure and protein behavior using giant plasma membrane vesicles. Nat Protoc 7, 1042–1051 (2012).

22. A. S. Klymchenko, R. Kreder, Fluorescent probes for lipid rafts: from model membranes to living cells. Chem Biol 21, 97–113 (2014).

23. B. B. Diaz-Rohrer, K. R. Levental, K. Simons, I. Levental, Membrane raft association is a determinant of plasma membrane localization. Proceedings of the National Academy of Sciences of the United States of America 111, 8500–8505 (2014).

24. I. Levental, D. Lingwood, M. Grzybek, U. Coskun, K. Simons, Palmitoylation regulates raft affinity for the majority of integral raft proteins. Proc Natl Acad Sci U S A 107, 22050–22054 (2010).

25. K. Raghunathan et al., Glycolipid Crosslinking Is Required for Cholera Toxin to Partition Into and Stabilize Ordered Domains. Biophysical journal 111, 2547–2550 (2016).

26. P. Sengupta, A. Hammond, D. Holowka, B. Baird, Structural determinants for partitioning of lipids and proteins between coexisting fluid phases in giant plasma membrane vesicles. Biochim Biophys Acta 1778, 20–32 (2008).

27. G. J. Snipes, U. Suter, A. A. Welcher, E. M. Shooter, Characterization of a novel peripheral nervous system myelin protein (PMP-22/SR13). J Cell Biol 117, 225–238 (1992).

28. J. Li, B. Parker, C. Martyn, C. Natarajan, J. Guo, The PMP22 gene and its related diseases. Mol Neurobiol 47, 673–698 (2013).

29. B. W. van Paassen et al., PMP22 related neuropathies: Charcot-Marie-Tooth disease type 1A and Hereditary Neuropathy with liability to Pressure Palsies. Orphanet J Rare Dis 9, 38 (2014).

30. S. Lee et al., PMP22 is critical for actin-mediated cellular functions and for establishing lipid rafts. J Neurosci 34, 16140–16152 (2014).

31. A. M. Jetten, U. Suter, The peripheral myelin protein 22 and epithelial membrane protein family. Prog Nucleic Acid Res Mol Biol 64, 97–129 (2000).

32. Y. Zhou et al., PMP22 regulates cholesterol trafficking and ABCA1-mediated cholesterol efflux. J Neurosci, (2019).

33. G. Saher et al., High cholesterol level is essential for myelin membrane growth. Nat Neurosci 8, 468–475 (2005).

34. G. Gopalakrishnan et al., Lipidome and proteome map of myelin membranes. J Neurosci Res 91, 321–334 (2013).

35. U. Suter, S. S. Scherer, Disease mechanisms in inherited neuropathies. Nat Rev Neurosci 4, 714–726 (2003).

36. S. Larrouquere-Regnier, F. Boiron, D. Darriet, C. Cassagne, J. M. Bourre, Lipid composition of sciatic nerve from dysmyelinating trembler mouse. Neurosci Lett 15, 135–139 (1979).

37. Y. Zhou et al., A neutral lipid-enriched diet improves myelination and alleviates peripheral nerve pathology in neuropathic mice. Exp Neurol 321, 113031 (2019).

38. R. Fledrich et al., Targeting myelin lipid metabolism as a potential therapeutic strategy in a model of CMT1A neuropathy. Nat Commun 9, 3025 (2018).

39. B. Hasse, F. Bosse, H. W. Muller, Proteins of peripheral myelin are associated with glycosphingolipid/cholesterol-enriched membranes. J Neurosci Res 69, 227–232 (2002).

40. N. Liu, J. Yamauchi, E. M. Shooter, Recessive, but not dominant, mutations in peripheral myelin protein 22 gene show unique patterns of aggregation and intracellular trafficking. Neurobiol Dis 17, 300–309 (2004).

41. J. P. Schlebach et al., Conformational Stability and Pathogenic Misfolding of the Integral Membrane Protein PMP22. J Am Chem Soc 137, 8758–8768 (2015).

42. M. C. Ryan, L. Notterpek, A. R. Tobler, N. Liu, E. M. Shooter, Role of the peripheral myelin protein 22 N-linked glycan in oligomer stability. J Neurochem 75, 1465–1474 (2000).

43. D. D’Urso, P. Ehrhardt, H. W. Muller, Peripheral myelin protein 22 and protein zero: a novel association in peripheral nervous system myelin. J Neurosci 19, 3396–3403 (1999).

44. S. J. Zoltewicz et al., The palmitoylation state of PMP22 modulates epithelial cell morphology and migration. ASN Neuro 4, 409–421 (2012).

45. Y. Song, A. K. Kenworthy, C. R. Sanders, Cholesterol as a co-solvent and a ligand for membrane proteins. Protein Sci 23, 1–22 (2014).

46. J. Fantini, C. Di Scala, C. J. Baier, F. J. Barrantes, Molecular mechanisms of protein-cholesterol interactions in plasma membranes: Functional distinction between topological (tilted) and consensus (CARC/CRAC) domains. Chem Phys Lipids 199, 52–60 (2016).

47. C. Di Scala et al., Relevance of CARC and CRAC Cholesterol-Recognition Motifs in the Nicotinic Acetylcholine Receptor and Other Membrane-Bound Receptors. Curr Top Membr 80, 3–23 (2017).

48. G. C. Li, E. T. C. Forster-Benson, C. R. Sanders, Genetic intolerance analysis as a tool for protein science. Biochim Biophys Acta Biomembr, 183058 (2019).

49. M. Silk, S. Petrovski, D. B. Ascher, MTR-Viewer: identifying regions within genes under purifying selection. Nucleic Acids Res 47, W121–W126 (2019).

50. K. F. Mittendorf, B. M. Kroncke, J. Meiler, C. R. Sanders, The homology model of PMP22 suggests mutations resulting in peripheral neuropathy disrupt transmembrane helix packing. Biochemistry 53, 6139–6141 (2014).

51. M. Sakakura, A. Hadziselimovic, Z. Wang, K. L. Schey, C. R. Sanders, Structural basis for the Trembler-J phenotype of Charcot-Marie-Tooth disease. Structure 19, 1160–1169 (2011).

52. I. Levental, M. Grzybek, K. Simons, Raft domains of variable properties and compositions in plasma membrane vesicles. Proc Natl Acad Sci U S A 108, 11411–11416 (2011).

53. A. Mohamed, H. Robinson, P. J. Erramouspe, M. M. Hill, Advances and challenges in understanding the role of the lipid raft proteome in human health. Expert Rev Proteomics 15, 1053–1063 (2018).

54. S. Minogue, M. G. Waugh, Lipid rafts, microdomain heterogeneity and inter-organelle contacts: impacts on membrane preparation for proteomic studies. Biol Cell 104, 618–627 (2012).

55. Y. Z. Zheng, L. J. Foster, Contributions of quantitative proteomics to understanding membrane microdomains. J Lipid Res 50, 1976–1985 (2009).

56. H. J. Sharpe, T. J. Stevens, S. Munro, A comprehensive comparison of transmembrane domains reveals organelle-specific properties. Cell 142, 158–169 (2010).

57. J. P. Schlebach et al., Topologically Diverse Human Membrane Proteins Partition to Liquid-Disordered Domains in Phase-Separated Lipid Vesicles. Biochemistry 55, 985–988 (2016).

58. H. Shogomori et al., Palmitoylation and intracellular domain interactions both contribute to raft targeting of linker for activation of T cells. J Biol Chem 280, 18931–18942 (2005).

59. S. A. Johnson et al., Temperature-dependent phase behavior and protein partitioning in giant plasma membrane vesicles. Biochim Biophys Acta 1798, 1427–1435 (2010).

60. J. Podkalicka, A. Biernatowska, M. Majkowski, M. Grzybek, A. F. Sikorski, MPP1 as a Factor Regulating Phase Separation in Giant Plasma Membrane-Derived Vesicles. Biophys J 108, 2201–2211 (2015).

61. C. Raggi et al., Caveolin-1 Endows Order in Cholesterol-Rich Detergent Resistant Membranes. Biomolecules 9, (2019).

