## Supporting Information for "Peripheral Myelin Protein 22 Preferentially Partitions into Ordered Phase Membrane Domains"

**This file includes:**

**Figure S1** Example of $P_{ordered}$ calculation.

**Figure S2** Localization of PMP22 variants in fixed cells.

**Figure S3** Phase preference for tgLAT in GPMVs derived from HeLa or RSCs.

**Figure S4** A different membrane phase marker does not alter $P_{ordered}$ for PMP22.

**Figure S5** Uncut western blots from palmitoylation calculation.

**Figure S6** $T_{Misc}$ for tgLAT containing GPMVs.

**Figure S7** Raw data for calculation of the ordered phase domain sizes in GPMVs

**Supporting Figure Captions**

**Figure S1**. **Example $P_{ordered}$ calculation. (A)** Three independent lines are drawn through the GPMV that pass through both an ordered and disordered phase of the membrane using the DiIC12 channel to distinguish the phases. Scale bar, 10 $\mu$m. **(B)** Fluorescent line intensity scans for the DiIC12 channel (red) and PMP22 channel (green). Membrane phases are indicated. **(C)** $P_{ordered}$ calculation for each individual line, with the mean value and standard deviation (SD) of $P_{ordered}$ being reported. **(D)** Definition of membrane phase preference from the $P_{ordered}$ values.

**Figure S2**. **Localization of PMP22 variants in cells.** Fixed and permeabilized cells were imaged via confocal microscopy using a 40x objective and an optical zoom of 3-4X. Propidium Iodide (PI, red) was used to identify the nuclei of cells and PMP22 was identified immunochemically via the myc epitope (green). Scale bar, 10 $\mu$m.

**Figure S3**. **Phase preference for tgLAT in GPMVs derived from HeLa or RSCs. (A)** Representative tgLAT-containing GPMVs derived from either HeLa or RSCs. Scale bar, 5 $\mu$m. **(B)** Quantification of $P_{ordered}$ for tgLAT from three independent biological experiments.

**Figure S4. A different membrane phase marker does not alter $P_{ordered}$ for PMP22**. **(A)** Representative N41Q PMP22 (red) containing GPMVs stained with the ordered phase marker NBD-DSPE (green). Scale bar, 5 $\mu$m. **(B)** Quantification of $P_{ordered}$ for N41Q PMP22 using the NBD-DSPE phase marker from three independent biological experiments.

**Figure S5. Uncut Western Blots from Palmitoylation calculation. (A)** Uncut anti-myc western blot from **Figure 2C**. **(B)** Uncut anti-biotin western blot from **Figure 2C**.

**Figure S6**. **$T_{Misc}$ for tgLAT containing GPMVs.** Fraction of tgLAT-containing GPMVs showing phase separation at various temperatures from 12.5°C-32.5°C were calculated for three independent biological replicates and fit to a sigmoidal curve. >100 GPMVs were imaged at each temperature for each experiment. $T_{Misc}$ is calculated from the fit of the sigmoidal curve for each independent experiment.

**Figure S7**. **Raw data for calculation of the ordered phase domain sizes in GPMVs. (A)** The average radius of GPMVs obtained from cells transfected with an empty vector (MOCK) or PMP22 containing GPMVs plotted in pixel size. **(B)** The percentage of vesicles containing ordered membrane phases. **(C)** Summary of results for all measurements.

**Figure S1**

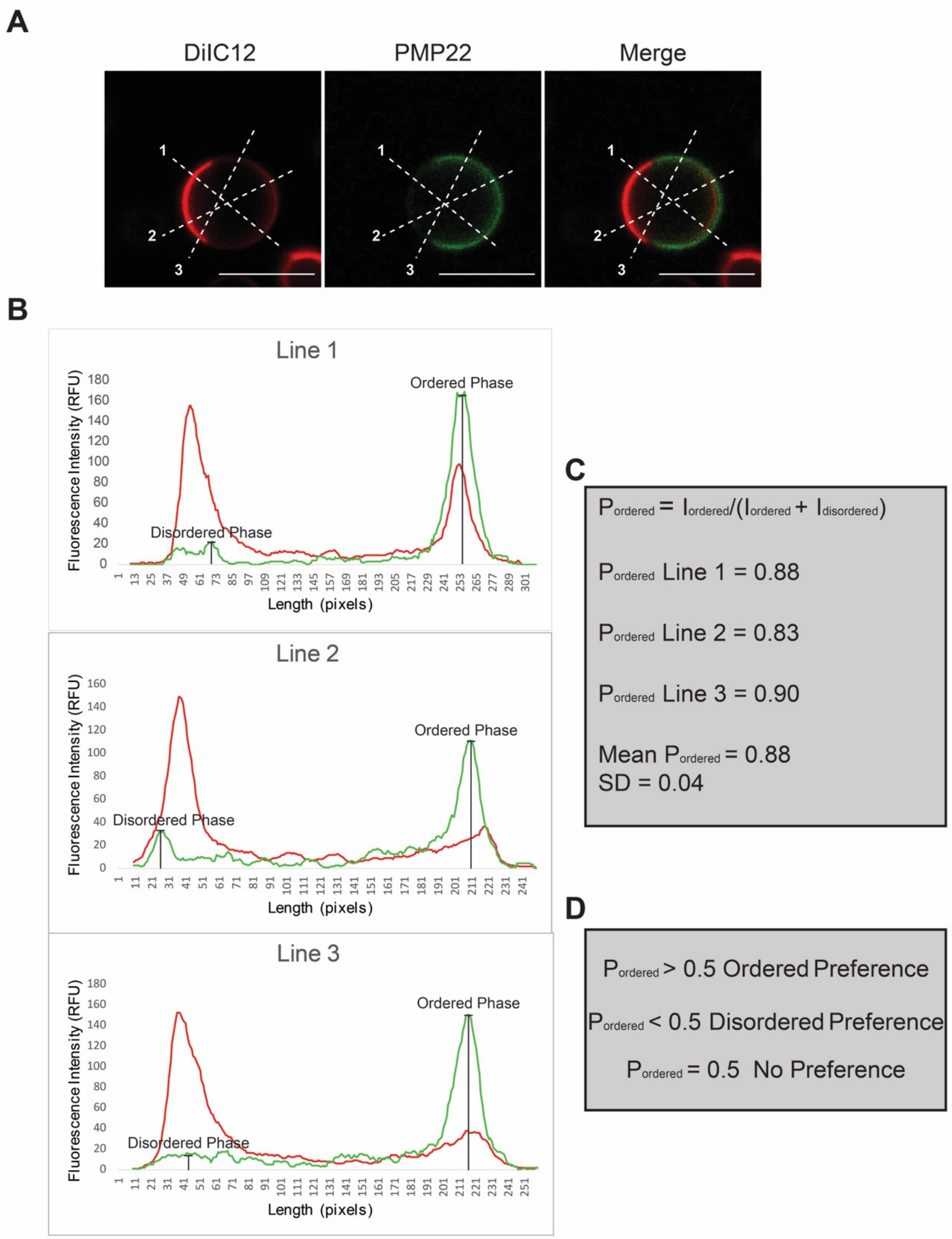


**Figure S2**

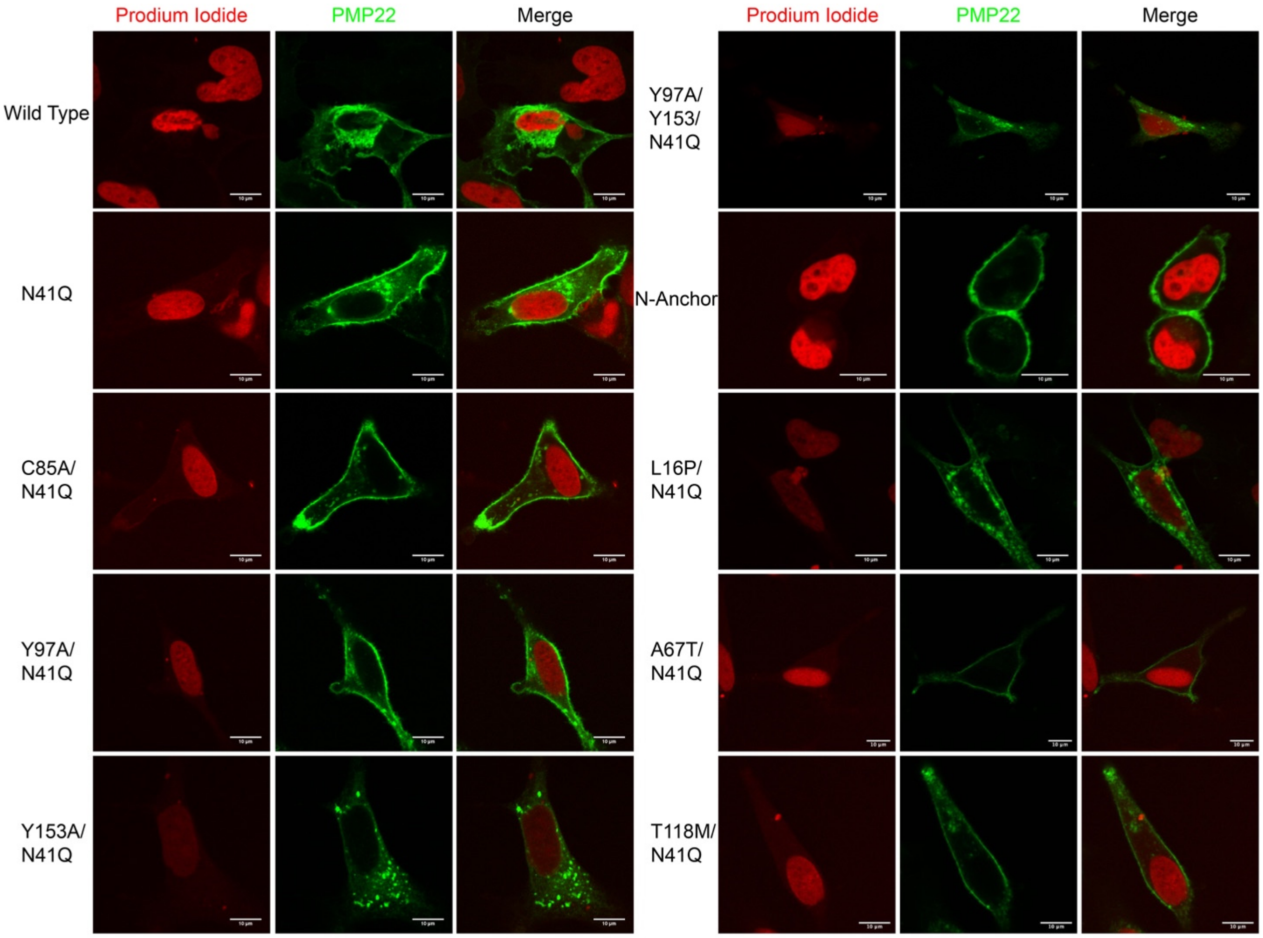

Prodium Iodide
PMP22
Merge
Prodium Iodide
PMP22
Merge
Wild Type
N41Q
C85A/
N41Q
Y97A/
N41Q
Y153A/
N41Q
Y97A/
Y153/
N41Q
N-Anchor
L16P/
N41Q
A67T/
N41Q
T118M/
N41Q


**Figure S3**

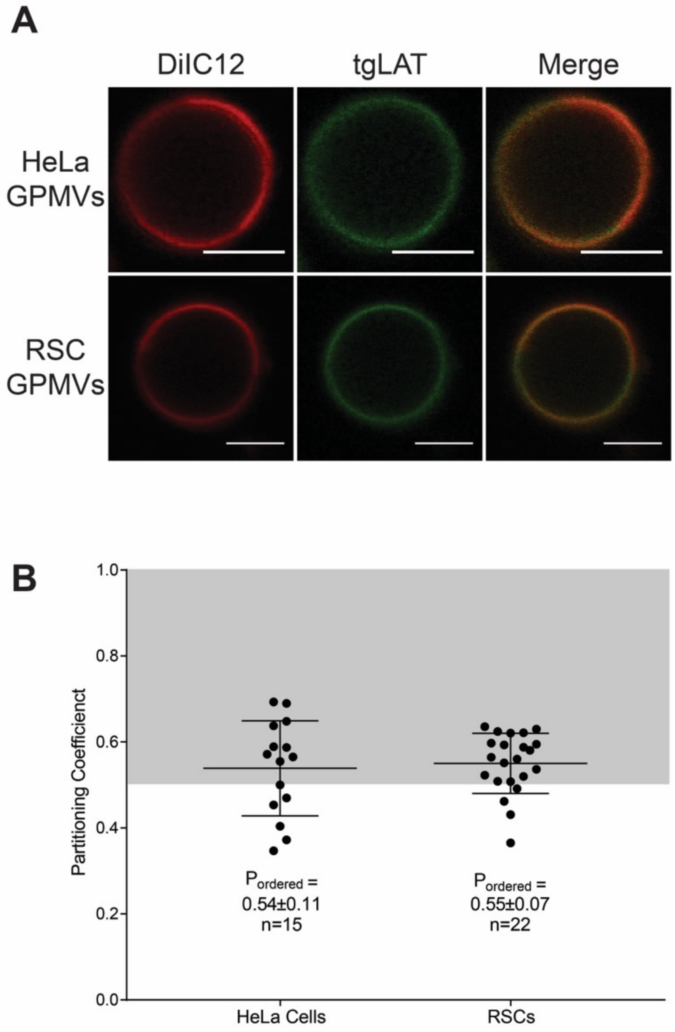
A
DiIC12
tgLAT
Merge
HeLa GPMVs
RSC GPMVs
B
Partitioning Coefficienct
P_ordered =
0.54±0.11
n=15
P_ordered =
0.55±0.07
n=22
HeLa Cells
RSCs
1.0
0.8
0.6
0.4
0.2
0.0

**Figure S4**

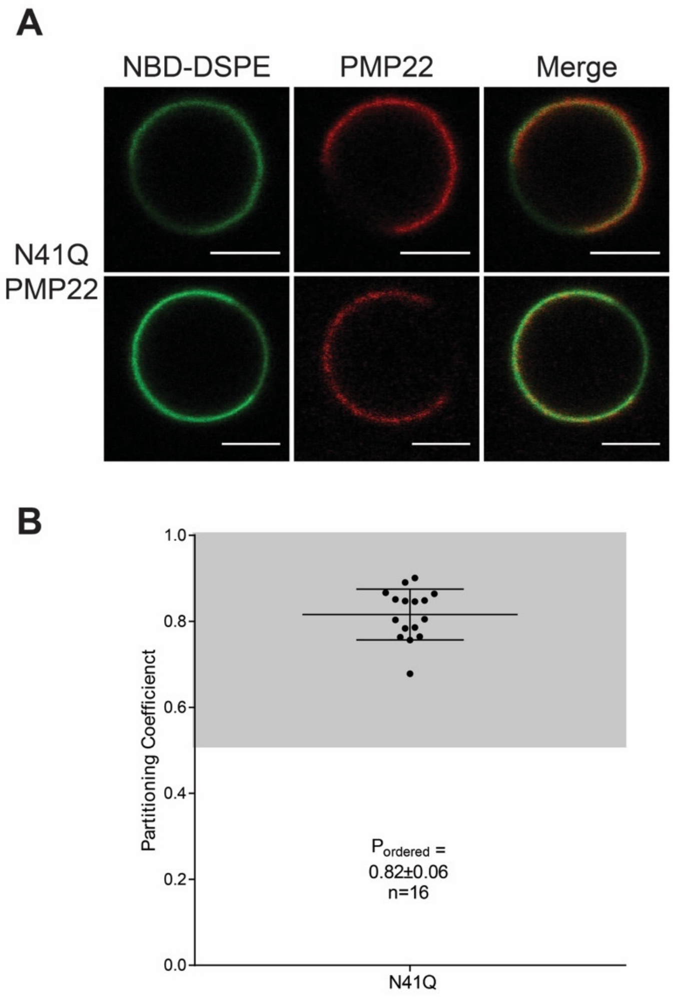

A
NBD-DSPE
PMP22
Merge
N41Q
PMP22
B
1.0
0.8
0.6
0.4
0.2
0.0
Partitioning Coefficient
P ordered =
0.82±0.06
n=16
N41Q

**Figure S5**

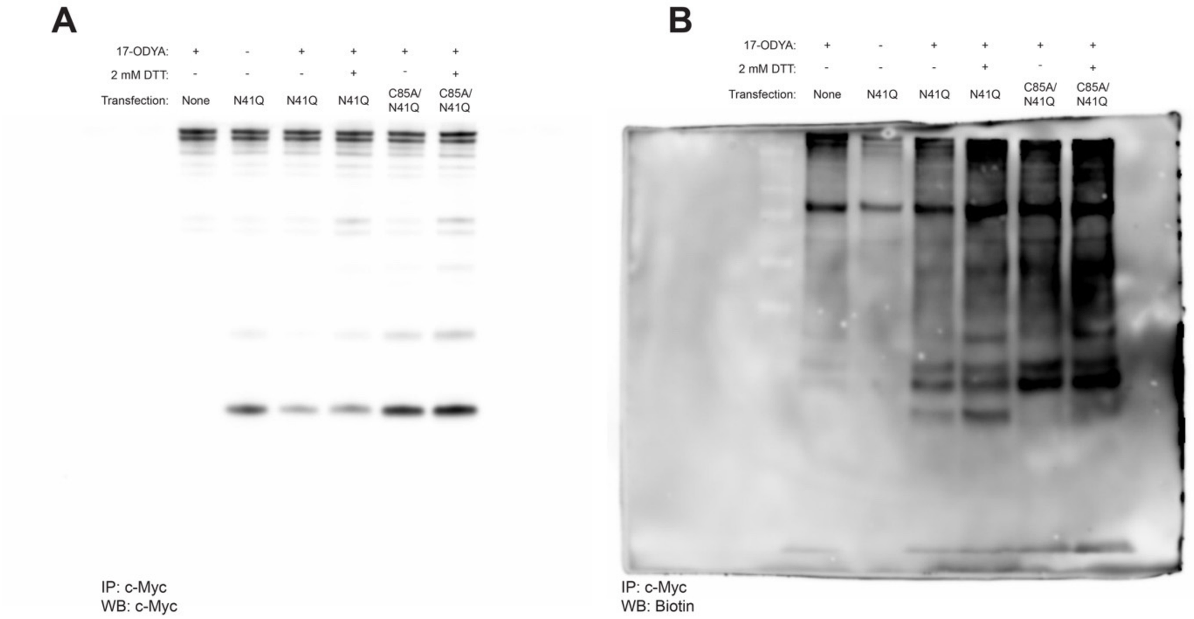
A
17-ODYA: + - + + + +
2 mM DTT: - - - + - +
Transfection: None N41Q N41Q N41Q C85A/ N41Q C85A/ N41Q
IP: c-Myc
WB: c-Myc
B
17-ODYA: + - + + + +
2 mM DTT: - - - + - +
Transfection: None N41Q N41Q N41Q C85A/ N41Q C85A/ N41Q
IP: c-Myc
WB: Biotin

**Figure S6**

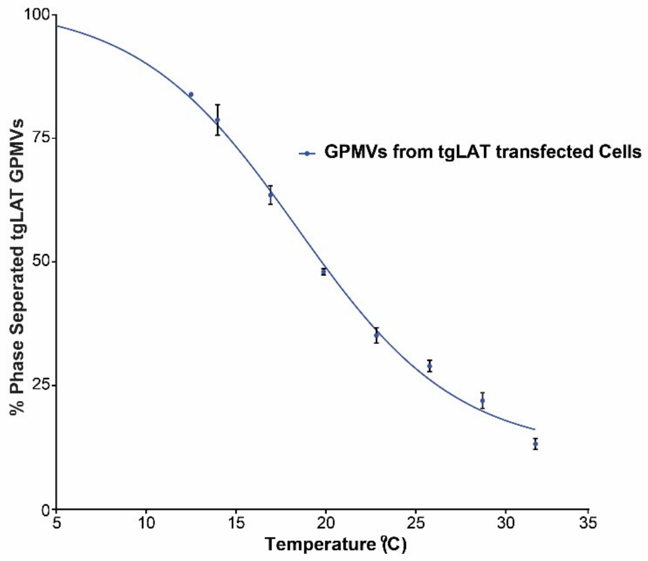

100
75
50
25
0
% Phase Seperated tgLAT GPMVs
5
10
15
20
25
30
35
Temperature (ºC)
GPMVs from tgLAT transfected Cells

### Figure S7

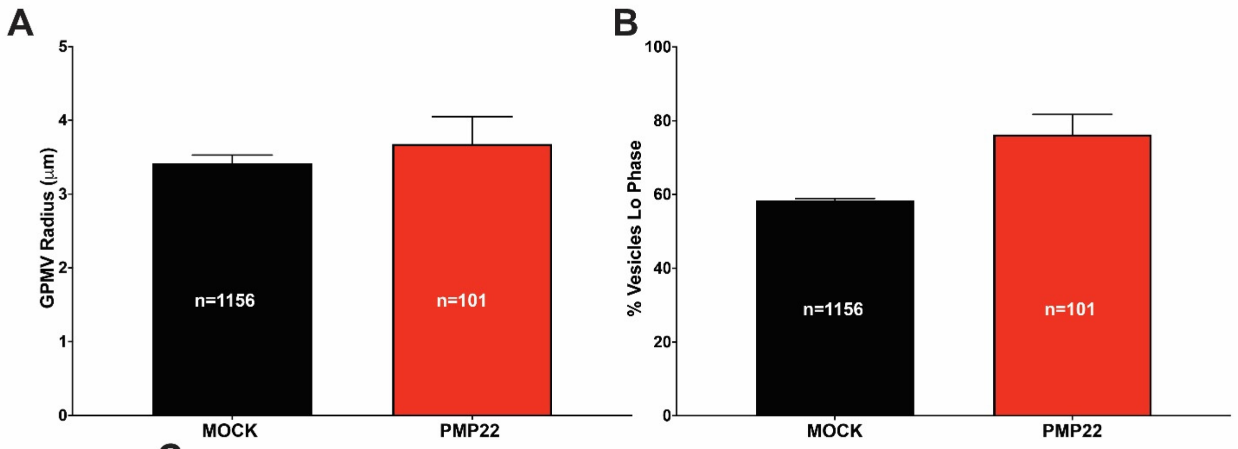


C

| | PMP22 | MOCK |
|---|---|---|
| Biological Replicates | 6 | 3 |
| Vesicles Measured | 101 | 1156 |
| Radius Vesicles (um) | 3.68±0.37 | 3.42±0.11 |
| Mean Circumference (um) | 23.13±2.31 | 21.48±0.70 |
| % of Vesicles Ordered Phase | 76.25±5.48 | 56.38±0.56 |
| Size of Ordered Domain (um) | 17.64±0.13 | 12.54±0.01 |
